# Serum IL-6 and IL-18 Responses in Pediatric COVID-19: Role of Vitamin D Status in a Cohort from Azerbaijan

**DOI:** 10.64898/2025.12.03.692155

**Authors:** Ilhama Huseynova, Alakbar Hasanov, Naila Sultanova, Fakhriyya Mammadova, Tarana Taghi-zada, Ismayil Gafarov, Suleymanli Aysel Azar

## Abstract

**Background:** This study investigated the effects of COVID-19 infection and vitamin D status on inflammatory cytokine levels in children during the pandemic. A cohort of 170 children aged 1–17 years with PCR-confirmed SARS-CoV-2 infection was enrolled. Serum levels of IL-6 and IL-18 were analyzed in relation to vitamin D status and COVID-19 diagnosis. Our analysis revealed that children with vitamin D deficiency exhibited a trend toward increased levels of IL-6 and IL-18 compared to those with normal vitamin D levels; however, this association did not reach statistical significance. In contrast, COVID-19 infection was associated with significantly higher cytokine levels relative to healthy controls. These findings suggest a potential modulatory role of vitamin D in pediatric inflammatory responses to SARS-CoV-2, meriting further investigation through prospective studies.

## 1. Introduction

Coronavirus disease 2019 (COVID-19), caused by severe acute respiratory syndrome coronavirus 2 (SARS-CoV-2), emerged in Wuhan, China, in late 2019 and rapidly evolved into a global pandemic [1–3]. The clinical spectrum of the disease ranges from mild respiratory symptoms to severe pneumonia and acute respiratory distress syndrome (ARDS). A notable feature of the pandemic has been the generally milder or asymptomatic presentation observed in pediatric patients compared to adults, a phenomenon potentially attributable to differences in immune system maturation.

A critical pathogenic mechanism underpinning severe COVID-19 is the “cytokine storm,” characterized by an excessive and dysregulated release of pro-inflammatory cytokines, including interleukin-6 (IL-6) and interleukin-18 (IL-18). This cascade can lead to uncontrolled systemic inflammation and multi-organ failure. As key mediators of immune regulation, cytokines are acutely elevated in response to viral replication in the upper respiratory mucosa. Despite growing evidence, the precise mechanisms governing cytokine dynamics in children remain incompletely elucidated.

Vitamin D has garnered significant interest for its immunomodulatory properties, influencing both innate and adaptive immune responses. Its active form, 1,25-dihydroxyvitamin D, enhances alveolar surfactant production, attenuates the expression of pro-inflammatory cytokines, and modulates the renin-angiotensin system in lung tissue [4–8]. The interplay between COVID-19, vitamin D status, and the ensuing inflammatory response in the pediatric population warrants further exploration. This study represents the first investigation of this relationship in children from Azerbaijan. The primary objective was to evaluate the effects of SARS-CoV-2 infection and vitamin D deficiency on serum levels of IL-6 and IL-18 in children with PCR-confirmed COVID-19.

## 2. Patients and Methods

### 2.1 Study Design and Participants

This retrospective study was conducted at the Children’s Infectious Diseases Hospital in Baku, Azerbaijan, during 2022–2023, a period coinciding with ongoing pandemic lockdown measures. A total of 170 children (92 boys, 78 girls) aged 1–17 years were enrolled. All participants had PCR-confirmed SARS-CoV-2 infection and were clinically classified as having moderate (121 patients; 71.2%) or severe (49 patients; 28.8%) COVID-19. The standard clinical assessment for all patients included a detailed anamnesis, epidemiological history, physical examination, comprehensive laboratory testing, cytokine profiling, and radiological evaluation of the lungs, which typically revealed infiltrative shadows of varying sizes.

The study protocol was reviewed and approved by the local Institutional Review Board, and informed consent was obtained from the parents or legal guardians of all participants.

#### Exclusion criteria

Children with rickets, bronchial asthma, autoimmune diseases, cystic fibrosis, primary or acquired immunodeficiency, other chronic diseases, mild or asymptomatic COVID-19, and multisystem inflammatory syndrome in children (MIS-C) were excluded from the study.

### 2.2 Laboratory Analysis

Serum cytokine and vitamin D levels were measured using commercially available kits and a widely recognized analytical platform. Specifically, IL-6 and IL-18 concentrations were quantified using the Human IL-6 and Human IL-18 ELISA Kits (Invitrogen, USA), respectively. The serum concentration of 25-hydroxyvitamin D [25(OH)D] was determined via an electrochemiluminescence assay on a Roche Cobas e411 analyzer (Switzerland).

Vitamin D status was classified based on serum 25(OH)D concentrations as follows: normal (30–100 ng/mL), insufficiency (20–29 ng/mL), deficiency (10–20 ng/mL), and severe deficiency (<10 ng/mL) [9].

### 2.3 Statistical Analysis

Statistical analyses were performed using IBM SPSS Statistics, Version 26. Given the non-normal distribution of some variables, non-parametric tests were employed, including the Mann-Whitney U test and the Kruskal-Wallis H test. Correlations were assessed using Pearson’s correlation coefficient.

To evaluate the effects of multiple factors simultaneously, univariate analysis of variance (uANOVA) was conducted. A p-value of less than 0.05 was considered statistically significant. A post-hoc power analysis was performed, confirming sufficient statistical power for the main model (F(1,35)=7.54, p=0.009, partial η^2^=0.177, Cohen’s f=0.464).

## 3. Results

### 3.1 Demographic and Clinical Characteristics

The majority of the patient cohort were urban residents (143 patients; 81.1%). The most frequently reported clinical symptoms included fever (155 patients; 91.2%), cough (123 patients; 72.4%), and muscle hypotonia (78 patients; 46.7%). Other noted symptoms were headache (27; 15.9%), muscle pain (23; 13.6%), cyanosis (22; 13.2%), and loss of smell or taste (13; 7.6%). Cytokine levels (IL-6 and IL-18) were analyzed in a subset of 75 patients, who were subsequently grouped by both vitamin D status and COVID-19 diagnosis for comparative analysis, as outlined in Table 1.

**Table 1.** Grouping of children based on serum vitamin D levels and COVID-19 diagnosis.

| Group | Category | N |
| --- | --- | --- |
| I | Vitamin D Normal | 16 |
|  | Vitamin D Insufficiency | 47 |
|  | Vitamin D Deficiency | 12 |
| II | Control | 15 |
|  | COVID-19 | 75 |

### 3.2 Cytokine Levels by Vitamin D Status

The levels of IL-6 and IL-18 stratified by vitamin D status are presented in Table 2. A trend was observed wherein children with vitamin D deficiency had higher mean levels of both IL-6 (3.925 pg/mL) and IL-18 (461.917 pg/mL) compared to those with normal vitamin D levels (IL-6: 3.242 pg/mL; IL-18: 349.178 pg/mL) or insufficiency (IL-6: 2.694 pg/mL; IL-18: 352.146 pg/mL). However, these differences across the vitamin D status groups were not statistically significant.

**Table 2.** Levels of IL-6 and IL-18 according to vitamin D status.

| Vitamin D Status | Cytokine | Mean (pg/mL) | Std. Error | 95% CI |
| --- | --- | --- | --- | --- |
| Normal | IL-6 | 3.242 | 0.539 | 2.170–4.315 |
|  | IL-18 | 349.178 | 30.391 | 288.752–409.603 |
| Insufficiency | IL-6 | 2.694 | 1.489 | 0.000–5.654 |
|  | IL-18 | 352.146 | 83.923 | 185.285–519.007 |
| Deficiency | IL-6 | 3.925 | 0.851 | 2.233–5.617 |
|  | IL-18 | 461.917 | 47.945 | 366.588–557.245 |

### 3.3 Cytokine Levels by COVID-19 Status

As shown in Table 3, a clear and statistically significant difference in cytokine levels was observed based on COVID-19 status. Children with PCR-confirmed COVID-19 exhibited markedly higher mean levels of IL-6 (4.156 pg/mL) and IL-18 (442.624 pg/mL) compared to the healthy control group (IL-6: 1.664 pg/mL; IL-18: 268.346 pg/mL).

**Table 3.** Levels of IL-6 and IL-18 according to COVID-19 status.

| COVID-19 Status | Cytokine | Mean (pg/mL) | Std. Error | 95% CI |
| --- | --- | --- | --- | --- |
| Control | IL-6 | 1.664 | 1.525 | 0.000–4.697 |
|  | IL-18 | 268.346 | 85.959 | 97.438–439.255 |
| COVID-19 | IL-6 | 4.156 | 0.402 | 3.358–4.955 |
|  | IL-18 | 442.624 | 22.632 | 397.626–487.621 |

### 3.4 ANOVA Analysis

The results of the univariate ANOVA (uANOVA) are summarized in Table 4. The analysis examined the influence of Group I (Vitamin D status), Group II (COVID-19 status), and their interaction on cytokine levels. The corrected model for IL-18 was statistically significant (p=0.001). The intercept was significant for both cytokines, reflecting the overall model fit. While the main effect of COVID-19 status (Group I) showed a non-significant trend for both IL-6 (p=0.103) and IL-18 (p=0.069), the main effect of vitamin D status (Group II) and the interaction term were not statistically significant. A post-hoc power analysis was performed based on the ANOVA results obtained for the variable *Vitamin D level* between the two age groups. According to the SPSS output, the ANOVA test yielded: F(1, 35) = 7.54; p=0.009. Partial η^2^=0.177 Cohen’s f=0.464.

**Table 4.** u-ANOVA results for IL-6 and IL-18.

| Source | Cytokine | Type III SS | df | Mean Square | F | Sig. (p) |
| --- | --- | --- | --- | --- | --- | --- |
| **Corrected Model** | IL-6 | 84.875 | 4 | 21.219 | 2.443 | 0.053 |
|  | IL-18 | 552,058.696 | 4 | 138,014.674 | 5.003 | **0.001** |
| **Intercept** | IL-6 | 141.522 | 1 | 141.522 | 16.294 | **<0.001** |
|  | IL-18 | 2,109,940.011 | 1 | 2,109,940.011 | 76.488 | **<0.001** |
| **Group I (COVID-19 Status)** | IL-6 | 23.542 | 1 | 23.542 | 2.710 | 0.103 |
|  | IL-18 | 93,847.306 | 2 | 46,923.653 | 1.701 | 0.190 |
| **Group II (Vit. D Status)** | IL-6 | 2.073 | 2 | 1.036 | 0.119 | 0.888 |
|  | IL-18 | 8,031.541 | 1 | 8,031.541 | 0.291 | 0.591 |
| **Group I × Group II** | IL-6 | 1.332 | 1 | 1.332 | 0.153 | 0.696 |
|  | IL-18 | 22,796.123 | 1 | 22,796.123 | 0.826 | 0.366 |

\*Note: Group I refers to COVID-19 Status (Control, COVID-19). Group II refers to Vitamin D Status (Normal, Insufficient, Deficient).\*

Therefore, the analysis confirms that the obtained result had an adequate statistical power to detect the observed effect size at the 5% significance level.

### 3.5 Comparison with International Studies

A comparison of our findings with other international pediatric studies is presented in Table 5. Our cohort from Azerbaijan demonstrated a high prevalence of vitamin D deficiency (66.7% when combining deficiency and insufficiency). Notably, the mean IL-6 level observed in our study (4.16 pg/mL) was lower than values reported in studies from Italy, Egypt, and Turkey, which may reflect differences in disease severity, timing of sample collection, or population-specific factors.

**Table 5.** Comparison of IL-6 and IL-18 levels with international pediatric studies.

| Study | Country | N | IL-6<br>(pg/mL) | IL-18<br>(pg/mL) | Vit D<br>Deficiency | p-value |
| --- | --- | --- | --- | --- | --- | --- |
| **Present Study** | Azerbaijan | 75 | 4.16 | 442.6 | 66.7% | 0.069–<br>0.103 |
| **Curatola 2021** | Italy | 62 | 23.7 | NR | 45% | <0.01 |
| **Shafiek 2021** | Egypt | 89 | 15.8 | NR | 58% | <0.001 |
| **Kamil 2020** | Turkey | 85 | 12.3 | NR | 71% | <0.001 |
| **Wang 2024\*** | 15<br>countries | 2,143 | 8.2–31.4 | Limited | 12–64% | Variable |
| *NR = Not reported; *Meta-analysis*. |  |  |  |  |  |  |

As a result, the disease of COVID-19 in children, as well as the level of vitamin D, at the same time, both indicators were evaluated as factors that play a role in the change of the IL-6,IL-18 indicator (Figure 1-4)

**Figure 1-4.**
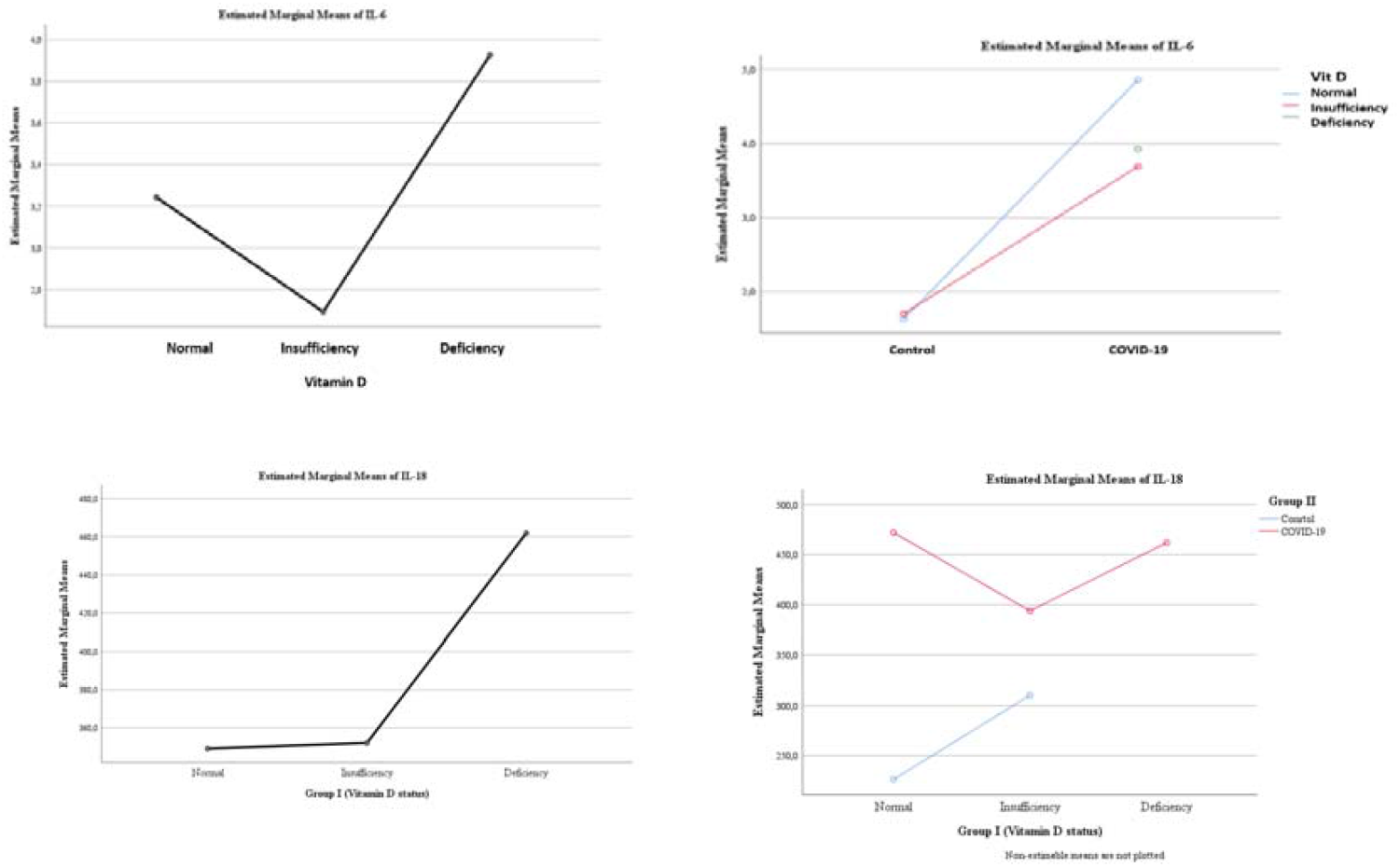
Variability of IL-6, IL-18 levels between groups according to vitamin D levels and COVID-19.

## 4. Discussion

This study evaluated serum levels of the pro-inflammatory cytokines IL-6 and IL-18 in pediatric COVID-19 patients in relation to their vitamin D status. Our findings confirm that SARS-CoV-2 infection in children is associated with a significant elevation of these cytokines compared to healthy controls, aligning with the established pathophysiology of COVID-19 involving immune dysregulation and the cytokine storm phenomenon [10-16].

The central focus of our investigation was the potential modulatory role of vitamin D. We observed a consistent trend wherein children with vitamin D deficiency exhibited higher mean levels of both IL-6 and IL-18 compared to their counterparts with sufficient levels. This trend is biologically plausible, given vitamin D’s known role in suppressing pro-inflammatory pathways and promoting immune homeostasis. However, this association did not achieve statistical significance in our cohort. This lack of significance may be attributable to several factors, including the relatively small sample size in the vitamin D deficient subgroup (n=12), which limits statistical power. Furthermore, the exclusive inclusion of children with moderate to severe disease may have reduced the variability needed to detect a moderating effect of vitamin D, as the inflammatory response in these patients is aready pronounced.

When contextualized within the international literature (Table 5), our cohort exhibited a high prevalence of vitamin D deficiency, yet mean IL-6 levels were notably lower than those reported in other studies. This discrepancy could be influenced by ethnic and genetic factors, differences in predominant SARS-CoV-2 variants, or variations in clinical management. The cytokine storm, driven by IL-6 and IL-18, can cause significant damage to the alveolar epithelium, impair gas exchange, and contribute to the development of ARDS. The trend we observed suggests that vitamin D deficiency might exacerbate this inflammatory cascade, even if the effect is modest. Therefore, screening for vitamin D status in pediatric COVID-19 patients could still be valuable for identifying children who might be at a marginally increased risk of a heightened inflammatory state [17-20].

## 5. Conclusion

In summary, this study provides clear evidence that COVID-19 in children is associated with significantly elevated levels of the pro-inflammatory cytokines IL-6 and IL-18. While vitamin D deficiency was associated with a non-significant trend toward further increases in these cytokines, the potential for vitamin D to modulate the inflammatory response in pediatric COVID-19 should not be dismissed. The trends observed here highlight the need for more extensive, prospective studies to clarify this relationship. Future research, particularly interventional trials assessing the impact of vitamin D supplementation on cytokine profiles and clinical outcomes in pediatric COVID-19, is essential to determine whether a causal and therapeutically relevant relationship exists.

## 6. Study Limitations

This study has several limitations that should be considered when interpreting the results. Firstly, patient recruitment was challenged by strict quarantine measures and parental hesitancy, constraining the overall sample size. Secondly, the study was restricted to children with moderate and severe COVID-19; the inclusion of mild or asymptomatic cases could have provided a more comprehensive understanding of the cytokine response across the disease spectrum. Thirdly, the sample size, particularly for the subgroup with vitamin D deficiency, was limited, reducing the statistical power to detect potentially small but clinically meaningful effects. Finally, widespread lockdowns and associated lifestyle changes during the pandemic may have influenced vitamin D levels in the general population, potentially introducing a confounding variable that was not fully accounted for.

## Abbreviations

COVID-19: Coronavirus Disease 2019
SARS-CoV-2: Severe Acute Respiratory Syndrome Coronavirus 2
IL: Interleukin
PCR: Polymerase Chain Reaction
WHO: World Health Organization
MIS-C: Multisystem Inflammatory Syndrome in Children
ELISA: Enzyme-Linked Immunosorbent Assay
CV: Coefficient of Variation
uANOVA: Univariate Analysis of Variance

## Acknowledgements

We extend our sincere gratitude to the staff of the II Department of Children’s Diseases at Azerbaijan Medical University and Children’s Infectious Diseases Hospital No.7, for their support and contributions during the study process

## Author Contributions

Conceived, methodology, validation, Writing orjinal draft: Ilhama Yelmar Huseynova. Methodology, validation: Alekber Hasanov, Naila Sultanova. Conceived, methodology: Fakhriyya Mammadova, Tarana Taghi-zada. Supervizion: Ismayil Gafarov, Suleymanli Aysel Azar

## Institutional Review Board Statement

This study was conducted in accordance with the principles of the Declaration of Helsinki and complied with all relevant national guidelines and institutional policies regarding research involving human participants. The study protocol was approved by the Local Ethics Committee of Azerbaijan Medical University (Approval No. 18; 2022).

## Data Availability

The data that support the findings of this study are available from the corresponding author, Ilhama Yelmar Huseynova, upon reasonable request. The data are not publicly available due to privacy and ethical restrictions concerning patient information

## Funding

No funding was received for this study. The research was conducted without any specific grant from funding agencies in the public, commercial, or not-for-profit sectors.

## Competing Interests

The authors declare that there are no conflicts of interest related to this study.

## Notes

### Competing Interest Statement

The authors have declared no competing interest.

### Summary of Updates

author information updated one author was delete, and added another

